# Tolerance induction in memory CD4 T cells is partial and reversible

**DOI:** 10.1101/2020.05.25.114785

**Authors:** Joshua I Gray, Shaima Al-Khabouri, Fraser Morton, Eric T Clambey, Laurent Gapin, Jennifer L Matsuda, John W Kappler, Philippa Marrack, Paul Garside, Thomas D Otto, Megan KL MacLeod

**Author notes:** Columbia Center for Translational Immunology, Columbia University, New York NY, USA 10032.

## Abstract

Memory T cells respond rapidly in part because they are less reliant on heightened levels of costimulatory molecules. This presents challenges to silencing memory T cells in tolerance strategies for autoimmunity or allergy. We find that memory CD4 T cells generated by infection or immunisation survive secondary activation with antigen delivered without adjuvant, regardless of their location in secondary lymphoid organs or peripheral tissues. These cells were, however, functionally altered following a tertiary immunisation with antigen and adjuvant, proliferating poorly but maintaining their ability to produce inflammatory cytokines. Transcriptional and cell cycle analysis of these memory CD4 T cells suggest they are unable to commit fully to cell division potentially because of low expression of DNA repair enzymes. In contrast, these memory CD4 T cells could proliferate following tertiary reactivation by viral re-infection. These data suggest that tolerance induction in memory CD4 T cells is partial and can be reversed.

## Introduction

Memory CD4 T cells play central roles in enhancing immune protection against pathogens the host has previously encountered(Jaigirdar and MacLeod, 2015). However, activated and memory CD4 T cells also contribute to disease processes in chronic inflammatory conditions, including rheumatoid arthritis and multiple sclerosis(Cope et al., 2007, McGinley et al., 2018, Raphael et al., 2020). Most current treatments for these conditions require continued use of drugs that dampen or deplete immune mediators or cells. A cure for these diseases will require deletion or retraining of the CD4 T cells that contribute to pathology.

Antigen-specific tolerance strategies have been used for many years to treat allergies and there are ongoing trials in autoimmune patients(Gunawardana and Durham, 2018, Pearson et al., 2017, Rayner and Isaacs, 2018, Serra and Santamaria, 2019) The underlying rationale for these strategies is based on our knowledge of tolerance induction in T cells, mainly developed from experiments examining TCR-activation of naïve CD4 T cells in the absence of costimulatory and inflammatory signals(Greenwald et al., 2005, Miller et al., 2007, Nurieva et al., 2011). Much less is known about the consequences of activating memory CD4 T cells through their TCR alone. Memory CD4 T cells can respond more quickly to a secondary challenge because, in part, they are less reliant on heightened level of costimulatory signals(Holzer et al., 2003, London et al., 2000, MacLeod et al., 2006). While this contributes to rapid pathogen control, this presents significant hurdles for treatments that aim to induce antigen-specific tolerance in autoimmunity, allergy or transplantation(Hartigan et al., 2019, MacLeod and Anderton, 2015).

A deeper understanding of the functional and molecular consequences of activating memory CD4 T cells with TCR signals alone is required to surmount these hurdles. We recently demonstrated that memory CD4 T cells reactivated with antigen delivered in the absence of adjuvant return to the memory pool and survive longterm(David et al., 2014). However, tertiary reactivation led to a curtailed response. Here we address two outstanding questions: 1. whether the consequences of reactivating memory CD4 T cells with antigen alone are similar in lymphoid organs and peripheral tissues and 2. what the underlying cellular changes responsible for the curtailed tertiary response are. These are important questions as the pathology for many autoimmune and allergic conditions is present in peripheral tissues, and understanding the mechanisms of memory CD4 T cell tolerance is essential to improve treatments and monitor therapeutic success. Our data show that while memory CD4 T cell responses are altered following exposure to tolerogenic signals, their ability to respond and produce inflammatory cytokines is not permanently restrained.

## Results

### Antigen-specific memory CD4 T cells in lymphoid organs and in peripheral tissues respond to tolerogenic signals but fail to increase in number

To investigate the consequence of reactivating memory CD4 T cells with tolerogenic signals (antigen delivered without adjuvant), we needed to track antigen-specific CD4 T cells following secondary and tertiary reactivation. To achieve this, we generated memory CD4 T cells in lymphoid organs and peripheral tissues by infecting C57BL/6 mice with WSN Influenza A virus (IAV) intranasally (i.n.). We used MHC class II tetramers containing the immunodominant IAV peptide nucleoprotein (NP)_311-325_ to identify NP_311-325_-specific memory CD4 T cells in the spleen, lung draining mediastinal lymph node (MedLN) and lung (Supplementary Figure 1).

Previously(David et al., 2014), we delivered antigen intravenously (i.v.) as it is a well-established tolerogenic injection route(David et al., 2014, Jenkins and Schwartz, 1987, Liblau et al., 1996). As expected, i.v. injection of NP_311-325_ and the adjuvant PolyIC led to an increase of T cells within the spleen in mice previously infected with IAV. However, there was no increase in the numbers of antigen-specific CD4 T cells in the MedLN or lung (Supplementary Figure 2). These data suggested that the antigen failed to access the MedLN and lung and that this delivery route could not be used to address our questions.

In contrast, i.n. instillation of peptide in the presence (immunogenic) or absence (tolerogenic) of PolyIC led to reactivation of the memory CD4 T cells in all three organs (Figure 1A-B). Importantly, delivery of peptide in the absence of adjuvant i.n. to naïve animals led to functional tolerance of the antigen-specific CD4 T cell population, validating this injection route for assessment of memory CD4 T cell tolerance induction (Supplementary Figure 3).

**Figure 1:**
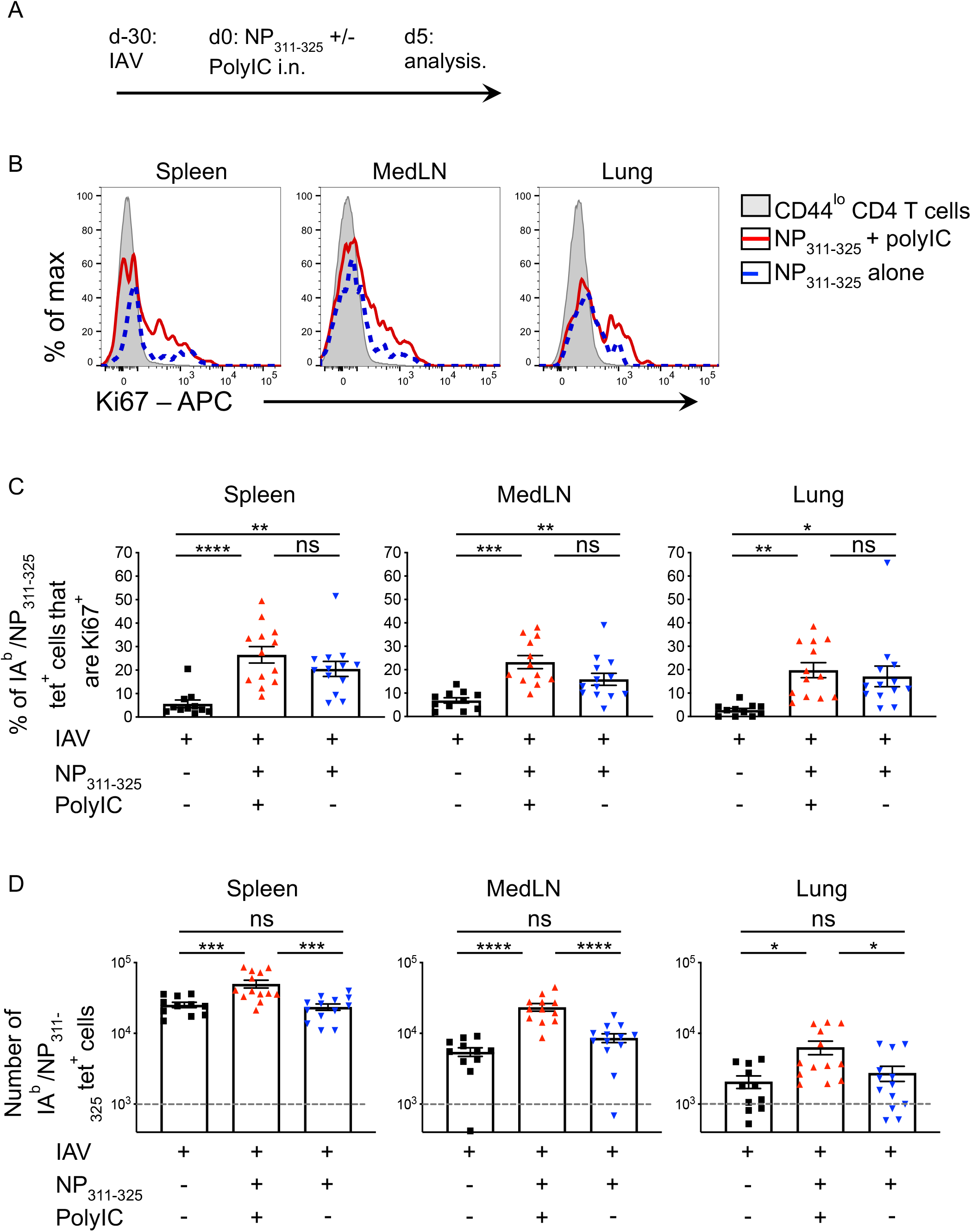
NP_311-325_-specific memory CD4 T cells reactivated with peptide delivered in the absence of adjuvant respond but fail to accumulate. C57BL/6 mice were infected with IAV on day −30. On day 0, some of these mice were immunised with NP_311-325_, +/−, PolyIC i.n. IA^b^/NP_311-325_ CD44^hi^ CD4 T cells were examined 5 days later in the spleen, MedLN, and lung (A) and their Ki67 expression (B, C) or their numbers determined (D). In C and D, each symbol represents one mouse and error bars are SEM. In D the grey dashed line represents the background staining in naïve animals. Data are combined from 3 experiments (3-5mice/experiment). Cells are gated as shown in Supplementary Figure 1. All statistics calculated using a one-way ANOVA with multiple comparisons; ns = not significant, * = <0.05, ** = <0.01, *** = <0.001, **** = <0.0001.

We first examined the immediate consequences of reactivating memory CD4 T cells with immunogenic or tolerogenic signals. Five days following instillation of NP_311-325_ peptide delivered with or without PolyIC, antigen-specific CD4 T cells showed evidence of activation via increased expression of the proliferation marker, Ki67, in all three organs (Figure 1B-C). Memory CD4 T cells reactivated in the presence of adjuvant increased in number as expected (Figure 1D). In contrast, there was no accumulation of memory CD4 T cells reactivated following the tolerogenic instillation of peptide alone. This suggests that while these cells entered the cell cycle, they either failed to complete mitosis or rapidly underwent cell death following proliferation.

### Memory CD4 T cells previously exposed to tolerogenic signals fail to expand upon subsequent reactivation despite entry into the cell cycle

To examine the longer term consequences of reactivating memory CD4 T cells with tolerogenic signals, we set up the experiment displayed in Figure 2A. Thirty days after infection with IAV, animals were given NP_311-325_ i.n. delivered with (immunogenic) or without (tolerogenic) PolyIC. After a further thirty days, we either examined the memory cells or performed a tertiary immunisation with NP_311-325_ conjugated to ovalbumin (OVA) protein delivered with the adjuvant alum.

**Figure 2:**
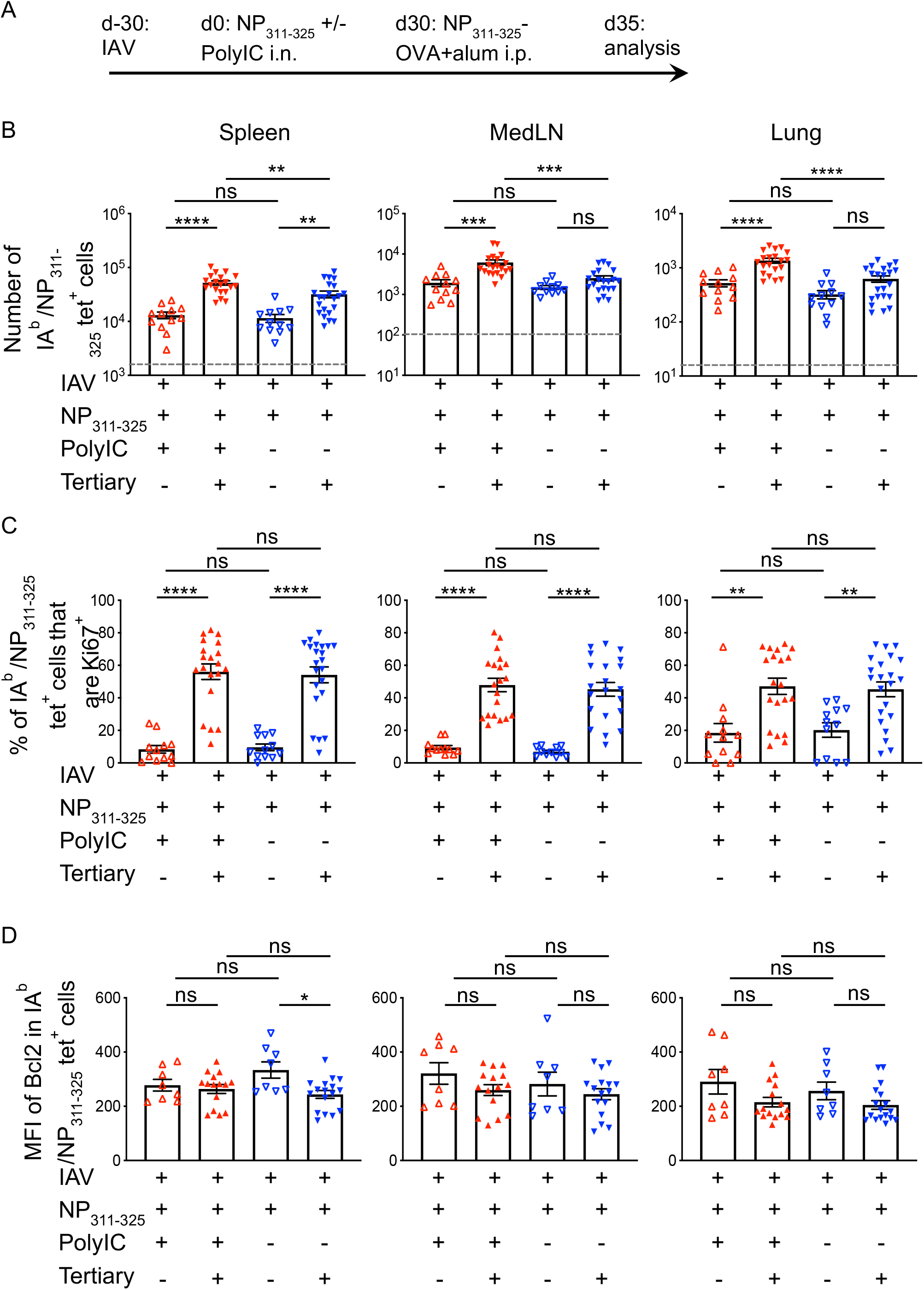
NP_311-325_-specific memory CD4 T cells previously reactivated with peptide without adjuvant fail to accumulate in the lung after tertiary reactivation. C57BL/6 mice were infected with IAV on day −30. On day 0, mice received NP_311-325_ +/− PolyIC and some of these mice were immunised i.p with NP-OVA with alum on day 30 (A). The numbers of IA^b^/NP_311-325_ CD44^hi^ CD4 T cells were examined 5 days later in the spleen, MedLN, and lung (B) and their expression of Ki67 (C) and Bcl2 (D) determined. Each symbol represents one mouse and error bars are SEM. In B, the grey dashed line represents the background staining in naïve animals. Data are combined from 4 experiments (4-8mice/experiment). All statistics calculated using a one-way ANOVA with multiple comparisons; ns = not significant, * = <0.05, ** = <0.01, *** = <0.001, **** = <0.0001.

Prior to tertiary reactivation, there were similar numbers of memory antigen-specific CD4 T cells in the two groups in each organ (Figure 2B). The antigen-specific CD4 T cells reactivated with an immunogenic secondary injection were able to mount a robust response in all organs upon tertiary reactivation. In contrast, the CD4 T cells previously exposed to tolerogenic signals expanded only slightly in the spleen and not at all in the MedLN and lung. There was, however, no difference in expression of Ki67 in the reactivated CD4 T cells (Figure 2C).

We also examined the expression of the pro-survival molecule Bcl2 to determine whether the memory CD4 T cells were more prone to apoptosis following tolerogenic activation (Figure 2D). However, there were no differences in the expression of Bcl2 between the two groups regardless of whether we examined the memory or the recalled cells in any of the three organs. This suggests increased apoptosis could not account for the poor accumulation of the tertiary reactivated memory CD4 T cells exposed to tolerogenic signals.

### Memory CD4 T cells reactivated with tolerogenic signals are not converted to regulatory T cells

Previous studies have shown that activation with antigen alone can lead to tolerance via the induction of FoxP3-expressing regulatory T cells(Zhang et al., 2013). We found few NP_311-325_-specific FoxP3^+^ cells at any timepoint examined (Figure 3). While there was a slightly increased percentage of antigen-specific FoxP3+ cells in the mice exposed to tolerogenic signals at one time point in the spleen, there were no differences in the number of FoxP3+ NP_311-325_-specific cells between the groups. This suggests that Treg conversion does not explain the poor accumulation of antigen-specific memory CD4 T cells previously exposed to tolerogenic signals.

**Figure 3:**
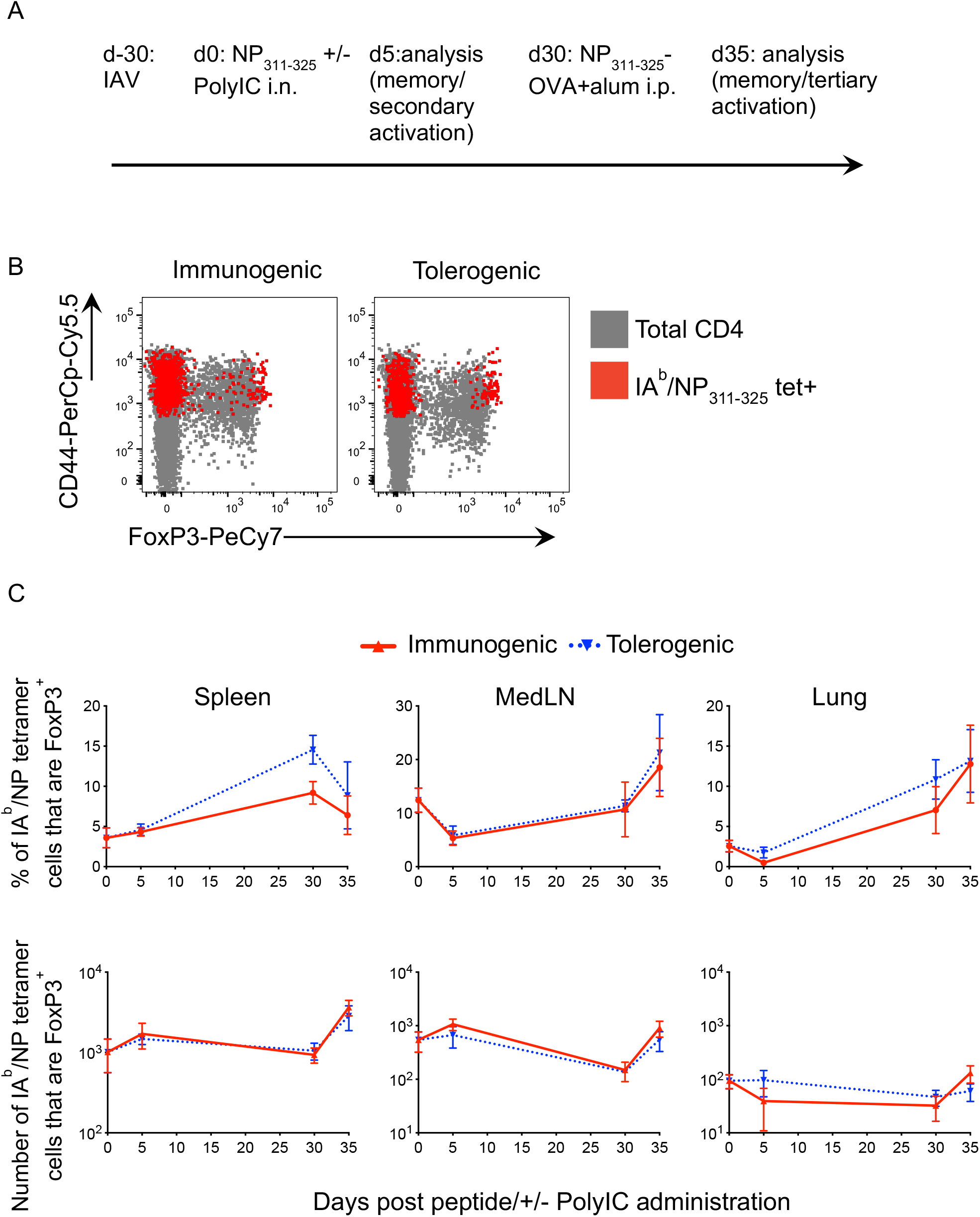
NP_311-325_-specific memory CD4 T cells exposed to soluble antigen in the absence of adjuvant are not converted to Tregs. C57BL/6 mice were infected with IAV on day −30. On day 0, mice received NP_311-325_ +/− PolyIC i.n. and some of these mice were immunised i.p with NP-OVA and alum on day 30 (A). The expression of FoxP3 by IA^b^/NP_311-325_ CD44^hi^ CD4 T cells was determined and shown in red and total CD4 T cells in grey in representative FACS plots (B) and the percentages and numbers of FoxP3 by IA^b^/NP_311-325_ CD44^hi^ CD4 T cells days 0, 5 and 35 shown in (C).The data are combined from 1 experiment/timepoint with 4-5 mice/group/timepoint. Each symbol represents the mean of between 4-5 mice and the error bars are SEM, *** = <0.001.

### Memory CD4 T cells exposed to tolerogenic signals display evidence of mitotic catastrophe following reactivation

Thus far our experiments indicated that memory CD4 T cells activated with tolerogenic signals can survive in the memory pool but accumulate poorly upon tertiary reactivation. To take an unbiased approach to investigate this failure, we performed transcriptomics analysis. For this we required a significant number of memory CD4 T cells that could be easily isolated for analysis. As identification of antigen-specific CD4 T cells by MHC tetramers requires ligation of the TCR by MHC molecules and the number of epitope specific cells are limited, we developed a novel triple transgenic reporter mouse TRACE (T cell Reporter of Activation and Cell Enumeration). We generated a transgenic animal in which the IL-2 promoter drives expression of rtTA. These animals were crossed to B6.Cg-Tg(tetO-cre)1Jaw/J mice and B6.129X1-Gt(ROSA)26SorTm(EYFP+)Cos mice. In these animals, T cells activated through the TCR when the animals are given doxycycline (Dox) become permanently EYFP+ (Supplementary Figure 4A).

Feeding of the Dox+ diet for one week was sufficient to induce a small population of EYFP+ cells, even in the absence of immunisation. However, delivery of OVA conjugated to 20μm polyethylene carboxylate beads in combination with the strong adjuvant combination of anti-CD40 and PolyIC(Kurche et al., 2010) drove an increased population of EYFP+ CD4 T cells above background (Supplementary Figure 4B,C). OVA was used for these studies as we required a protein containing multiple CD4 T cell epitopes which could be obtained free of contaminating microbial products, which would be present in recombinant IAV proteins. Moreover, we found that recombinant nucleoprotein had intrinsic adjuvant properties(Macleod et al., 2013).

To reactivate the memory CD4 T cells, we returned to systemic i.v. delivery of OVA protein as this is widely accepted as a consistent tolerogenic route(David et al., 2014, Jenkins and Schwartz, 1987, Liblau et al., 1996). We found that secondary immunisation with OVA in the TRACE mice led to anaphylactic shock, likely a consequence of anti-OVA antibodies. To avoid this, we sorted EYFP+ CD4 T cells at day 8 after immunisation and transferred these cells into naïve C57BL/6 animals that were injected with OVA or OVA and LPS i.v. after 22 days, 30 days since the cells were first primed. Thirty days following this, the recipient animals were immunised with OVA+alum and CD4+ EYFP+ cells isolated 5 days later for RNA-seq analysis (Figure 4A).

**Figure 4:**
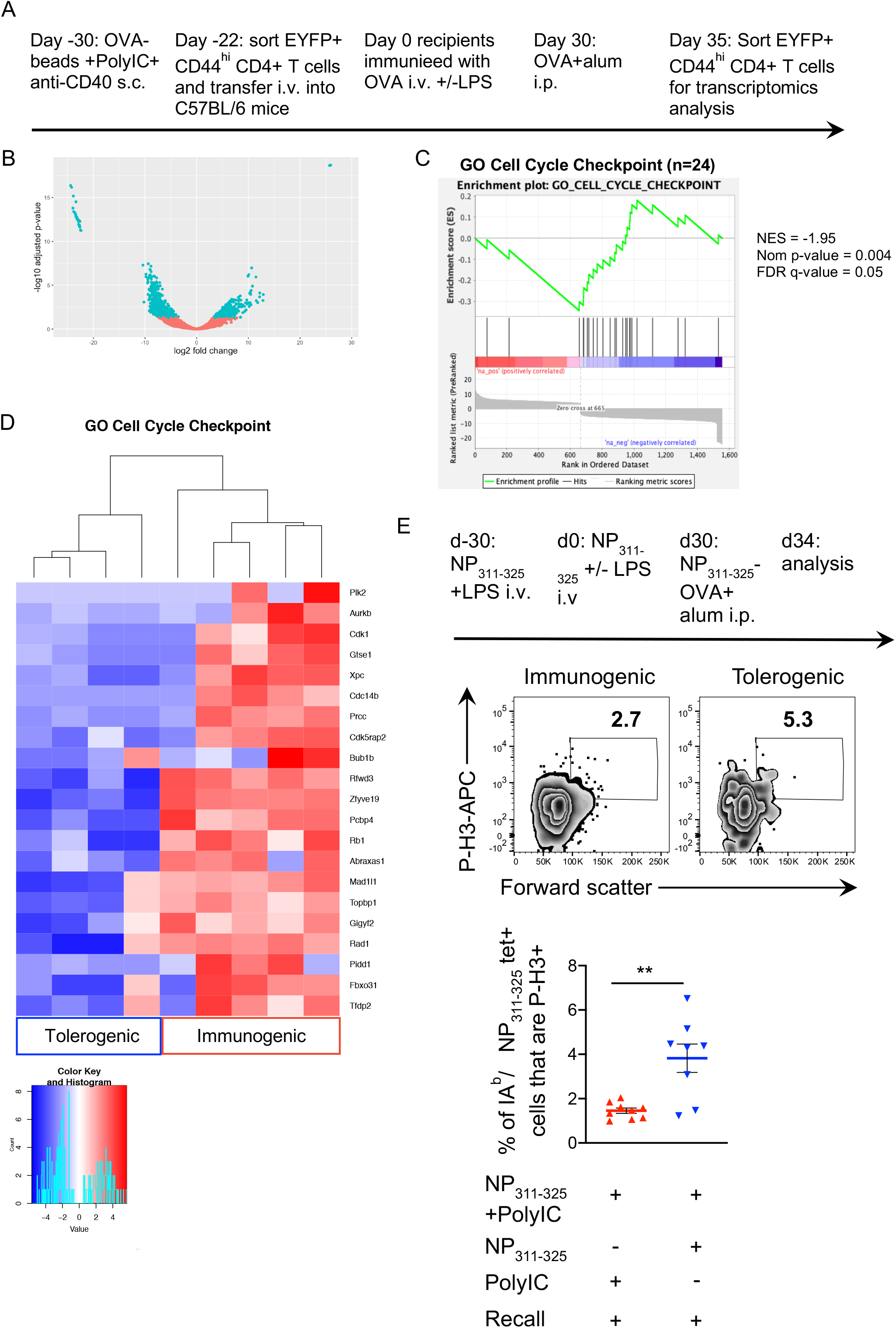
Transcriptomics and cell cycle analysis indicates that memory CD4 T cells exposed to tolerising signals undergo mitotic catastrophe following further reactivation in vivo. FACS sorted EYFP+ CD4 T cells from TRACE mice immunized with OVA+anti-CD40 and PolyIC were transferred into naïve C57BL/6 mice that were then immunised with OVA +/− LPS then re-immunised with OVA+alum i.p. 30 days later (A). EYFP+ CD4 T cells were FACS sorted after a further 5 days and RNA isolated for transcriptomic analysis. The DEGs are displayed in a volcano plot (B). GESA and heatmap show expression of DEGs contained within the GO term ‘Cell Cycle Checkpoints’ (GO: 0000075) (C). DEGs within the GO term and expressed at lower levels in the tolerogenic samples are displayed in a heatmap (D). C57BL/6 mice immunised with NP_311-325_ and LPS were injected with NP_311-325_ +/− LPS 30 days later and finally immunised after a further 30 days with NP-OVA+alum. 5 days later the percentages of forward scatter high IA^b^/NP_311-325_ tetramer+ cells that expressed phosphorylated Histone3 were examined (E). In (E) cells are gated as in supplementary Figure 1 and plots are concatenated from 4 mice per group. Data are combined from 2 experiment with 4-5 mice/group, error bars are SEM. Statistical analysis in B calculated by a T-test, ** = <0.01.

Gene expression from 5 individual mice in each experimental condition were analysed. One sample from the tolerogenic group was excluded as the number of EYFP+ CD4 T cells collected was 2.5-10fold higher than any of the other samples, suggesting an abnormal response or potential contamination during sorting. Of the differently expressed genes (DEGs), 898 were expressed at lower levels and 667 at higher levels in the tolerogenic group compared to the immunogenic group (Figure 4B). Analysis of DEGs expressed at higher levels in the tolerised samples failed to find consistent changes across all four samples. We, therefore, concentrated on DEGs that were expressed at lower levels in the tolerised samples. Gene ontogeny (Panther(Mi et al., 2019)) analysis of the biological processes associated with these DEGs indicated overrepresentation of gene products involved in ‘DNA-dependent DNA replication’, ‘spindle organisation’ and ‘cell cycle checkpoints’ (Table 1 and Supplementary Excel File 1).

**Table 1:**
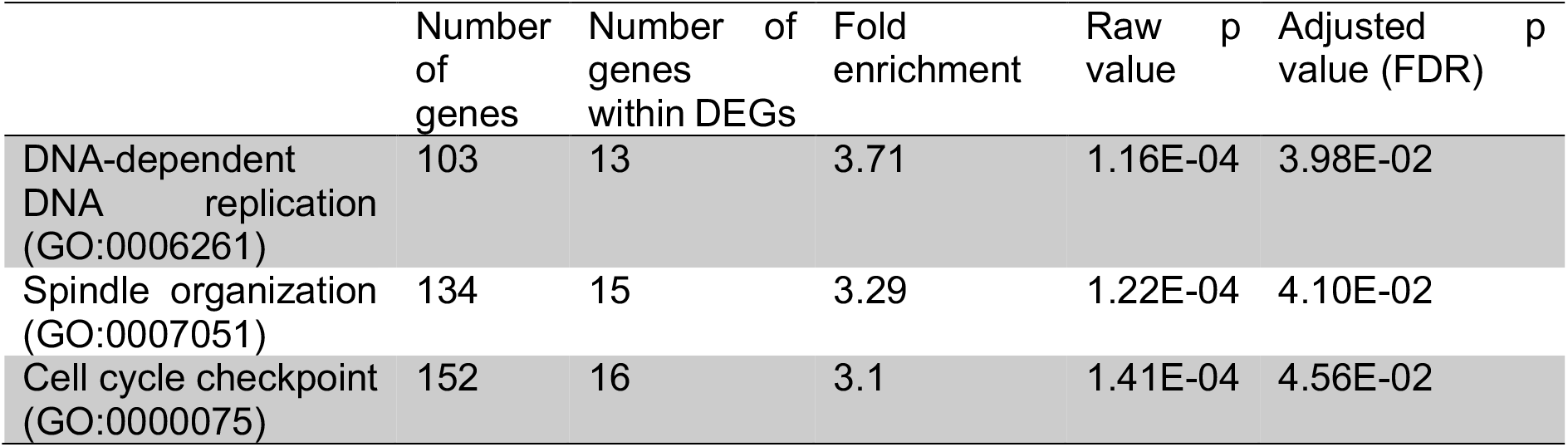
Gene over-representation analysis of DEGs expressed at lower levels in EYFP+ CD4 T cells in tolerogenic groups (top 3)

We performed gene set enrichment analysis and found that the DEGs were enriched for genes within the GO term, Cell Cycle Checkpoints (Figure 4C); DEGs within this GO term that were expressed at lower levels in the tolerogenic samples are displayed as a heatmap (Figure 4D). A number of these molecules play key roles at various stages of the cell cycle and in spindle formation and function. These genes include: the essential cyclin, Cdk1(Santamaria et al., 2007); Aurkb, a key component of the Chromosome Passenger Complex required for normal spindle assemble(Joukov and De Nicolo, 2018); Mad1l1 (also known as MAD1), a component of the spindle-assembly checkpoint(Hardwick and Murray, 1995, Musacchio, 2015); and Cdk5rap2 which plays a number of roles in spindle checkpoints(Lizarraga et al., 2010, Zhang et al., 2009).

Dysfunction of the spindle checkpoint is linked to death by mitotic catastrophe, a form of cell death that occurs when cells are unable to complete mitosis(Nitta et al., 2004, Shang et al., 2010). As the percentages of CD4 T cells that were Ki67+ after tertiary activation were equivalent regardless of whether or not they had been activated with immunogenic or tolerogenic signals (Figure 2C), these data suggests that memory CD4 T cells activated with tolerogenic signals can enter the cell cycle but fail to complete cell division following tertiary reactivation. To investigate this, we examined the proportion of reactivated memory antigen-specific T cells in mitosis reasoning that more CD4 T cells would be found in mitosis in reactivated cells previously exposed to tolerogenic signals as they would be ‘stuck’ in mitosis.

The percentages of re-activated antigen-specific T cells in mitosis were determined by expression of phosphorylated (p)Histone 3, present only during mitosis(Hans and Dimitrov, 2001). We focused on cells with increased forward scatter as cells increase in size during cell division(Bohmer et al., 2011). CD4 T cells were examined 4 days after tertiary reactivation of mice first immunised with NP_311-325_ and LPS, then reactivated with NP_311-325_ delivered with or without LPS and finally reactivated with NP-OVA+alum (Figure 4E). Very few antigen-specific T cells were positive for p-H3, but consistently more CD4 T cells were p-H3 positive in mice previously immunised with NP_311-325_ delivered without, than with, LPS (Figure 4E). This indicates that the cells in mice that received tolerogenic signals were more likely to be mitosis, suggestive of a failure to complete cell division.

Mitotic catastrophe often occurs in cells with DNA damage(Vakifahmetoglu et al., 2008). We, therefore, examined whether any DEGs were enriched in genes involved in GO term DNA repair. This was indeed the case (Figure 5A); the DEGs expressed at lower levels in the tolerised samples are shown as a heatmap (Figure 5B). The GSEA of the DEGs expressed at lower levels in the tolerogenic samples indicates that multiple genes contribute to this enrichment. These data suggest that tolerised memory CD4 T cells display poor repair of their DNA during reactivation-induced DNA synthesis and that this is a consequence of reduced expression of a number of different genes. This, coupled with the low expression of cell cycle checkpoint proteins, likely compromises their ability to commit to cell division.

**Figure 5:**
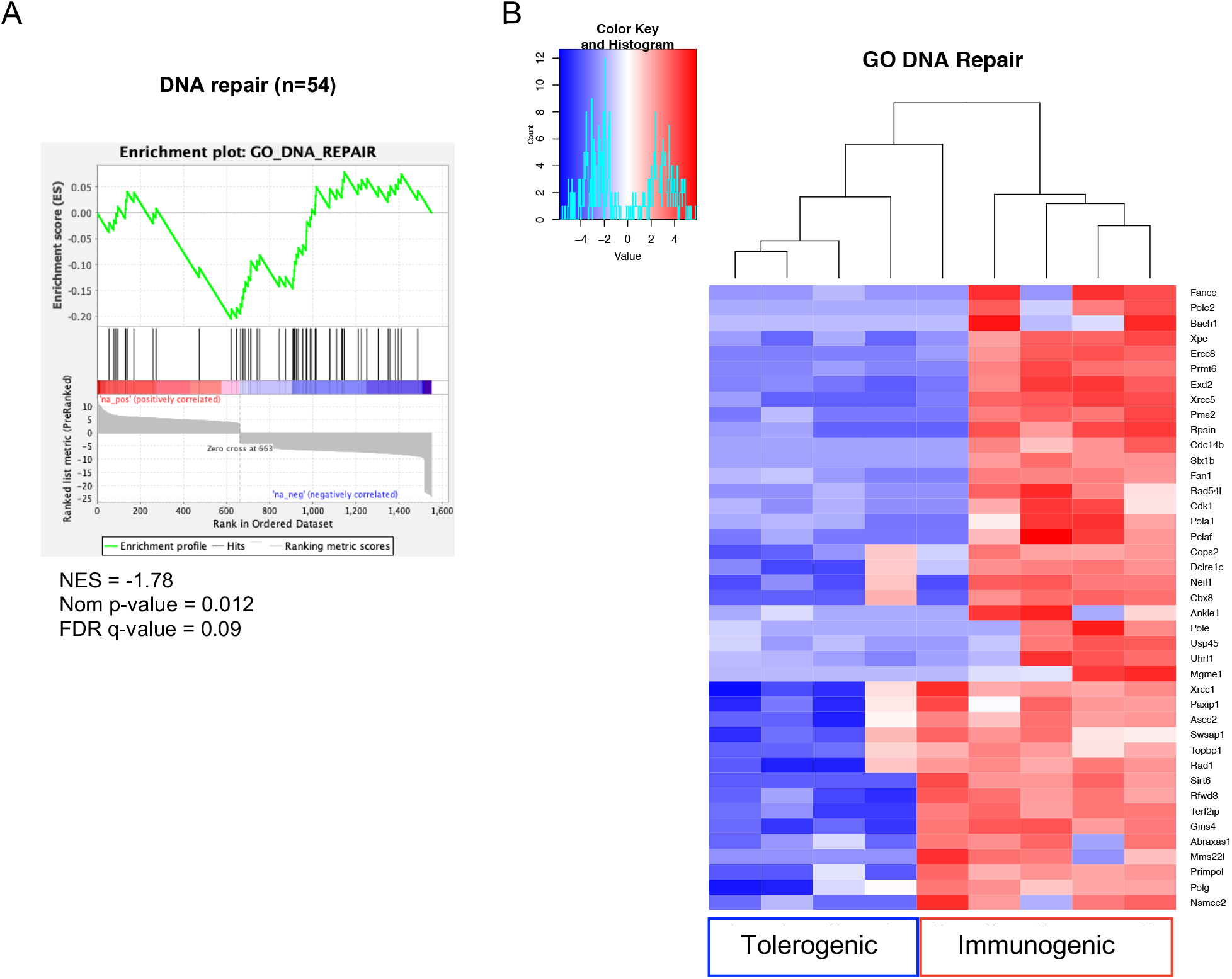
Transcriptomics analysis indicates that memory CD4 T cells exposed to tolerising signal have reduced expression of DNA repair enzymes. FACS sorted EYFP+ CD4 T cells from TRACE mice immunized with OVA+anti-CD40 and PolyIC were transferred into naïve C57BL/6 mice that were then immunised with OVA +/− LPS then re-immunised with OVA+alum i.p. 30 days later as in Figure 4. EYFP+ CD4 T cells were isolated by FACS after a further 5 days and RNA isolated for transcriptomic analysis. GSEA shows significant enrichment of the genes involved within the GO term ‘DNA repair’ (GO: 0006281) (A) and the DEGs expressed at lower levels in the tolerogenic samples are displayed in a heatmap (B).

### Memory CD4 T cells exposed to tolerogenic signals continue to produce cytokine but fail to provide accelerated help to primary responding B cells

Our data indicate that memory CD4 T cells reactivated with tolerogenic signals have impaired proliferative responses. We also wanted to determine whether these cells were impaired in other ways. To investigate this, we used the IAV infection model to generate sufficient cells in multiple organs to examine *ex vivo* cytokine production; cytokine responses are limited in antigen/adjuvant models(MacLeod et al., 2008).

Thirty days after mice were infected with IAV, they were injected with immunogenic or tolerogenic NP_311-325_ i.n. and then rested for 30 days (Figure 6A). Bone marrow dendritic cells loaded with NP_311-325_ were used to examine the *ex vivo* cytokine potential of the memory CD4 T cells and activated CD4 T cells from mice given a tertiary immunisation with NP_311-325_-OVA and alum delivered i.p. (Supplementary Figure 5).

**Figure 6:**
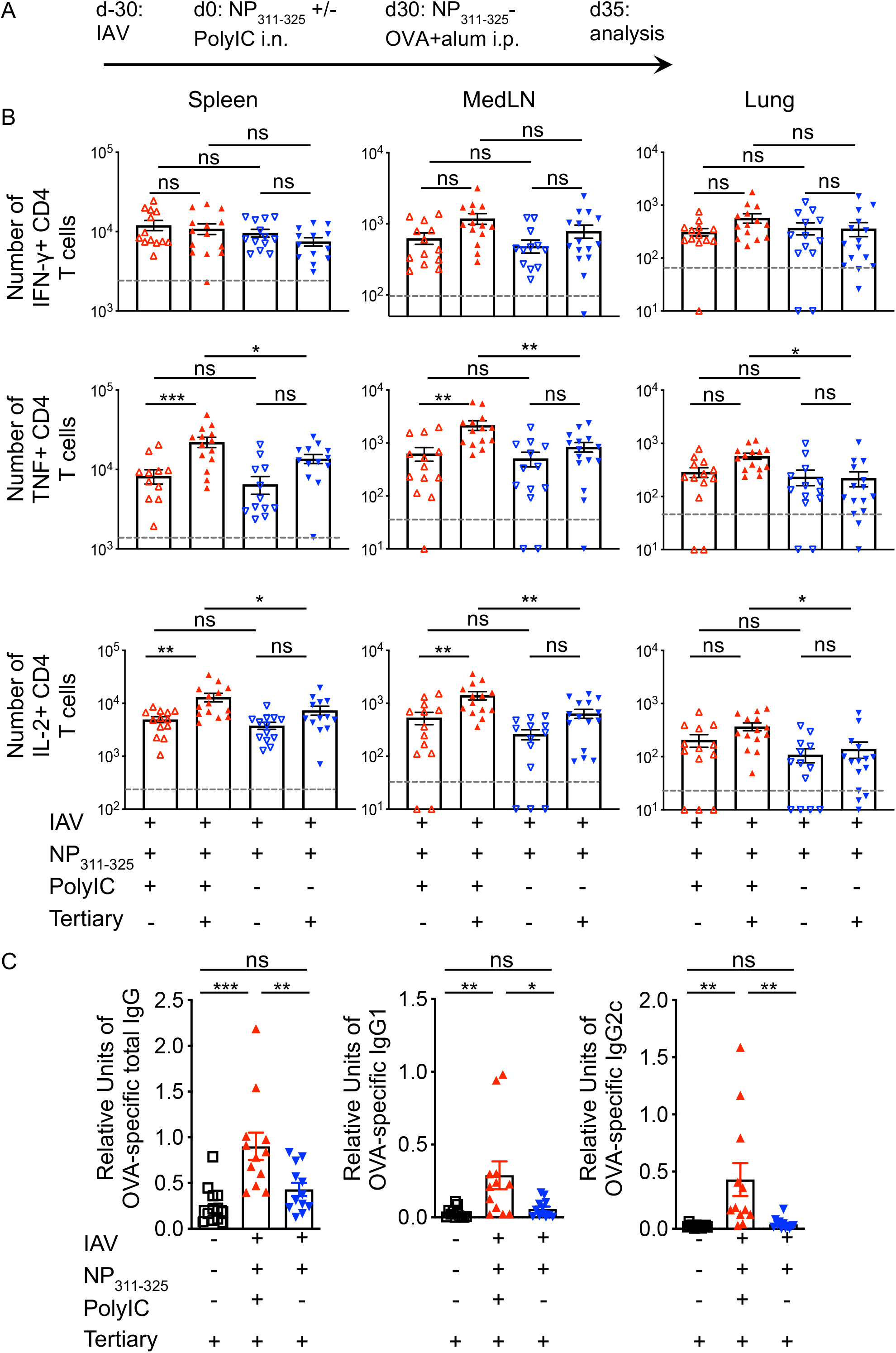
Activation of CD4 T cells with peptide in the absence of adjuvant does not affect CD4 T cell cytokine production but does prevent them providing accelerated help to B cells. C57BL/6 mice were infected with IAV on day −30. On day 0, mice received NP_311-325_ +/− PolyIC and some of these mice were immunised i.p with NP-OVA with alum on day 30 (A). On day 35, cells from the spleen, MedLN and lung were co-cultured with bmDCs loaded with NP_311-325_ for 6 hours in the presence of Golgi Plug and the number of IFN-γ, TNF and IL-2 producing CD44^hi^ CD4+ T cells examined (B). The levels of IgG, IgG1 and IgG2c anti-OVA antibodies in the serum was determined on day 5 (C). Each symbol represents one mouse and error bars are SEM. In B the grey dashed line represents the background staining in naïve animals. Data in B are combined from 2-3 experiments (3-5mice/experiment). Data in C are combined from 3 experiments with 4 mice/experiment. All statistics calculated using a one-way ANOVA with multiple comparisons; ns = not significant, * = <0.05, ** = <0.01, *** = <0.001, **** = <0.0001.

The numbers of IFN-γ, TNF or IL-2 producing antigen-specific memory CD4 T cells were equivalent in mice exposed to immunogenic or tolerogenic NP_311-325_ peptide 35 days previously (Figure 6B). Five days after reactivation with NP-OVA+alum, there was an increase of TNF and IL-2 producing cells in the spleen and the MedLN in mice previously exposed to NP_311-325_ and PolyIC. In contrast, there was no increase in the number of cytokine producing cells in mice previously exposed to tolerogenic NP_311-325_. In neither group did we see an increase in IFN-γ producing CD4 T cells. Together, these data suggest that, while exposure to tolerogenic signals affected accumulation of T cells, it did not prevent their ability to produce cytokines.

To investigate the functional responses of the T cells further, we examined their ability to provide accelerated help to primary responding OVA-specific B cells(David et al., 2014, MacLeod et al., 2011). We measured the levels of class-switched, OVA-specific antibody 5 days after the tertiary reactivation. As expected, primary responding mice had very little class-switched OVA-specific antibody and IAV-infected mice previously exposed to immunogenic signals had clearly detectable levels of OVA-specific immunoglobulin(MacLeod et al., 2011). In contrast, IAV-infected mice that had previously received tolerogenic signals failed to produce these antibodies at levels above primary immunised animals, demonstrating an impaired functional response (Figure 6C).

### Memory CD4 T cells exposed to tolerogenic signals expand following reactivation with influenza virus

Finally, we wanted to test whether the failure of CD4 T cells exposed to tolerogenic signals to accumulate could be rescued by reactivation with a more inflammatory stimulus. We therefore challenged IAV infected mice given immunogenic or tolerogenic signals with an heterosubtypic form of IAV, X31 (Figure 7A).

**Figure 7:**
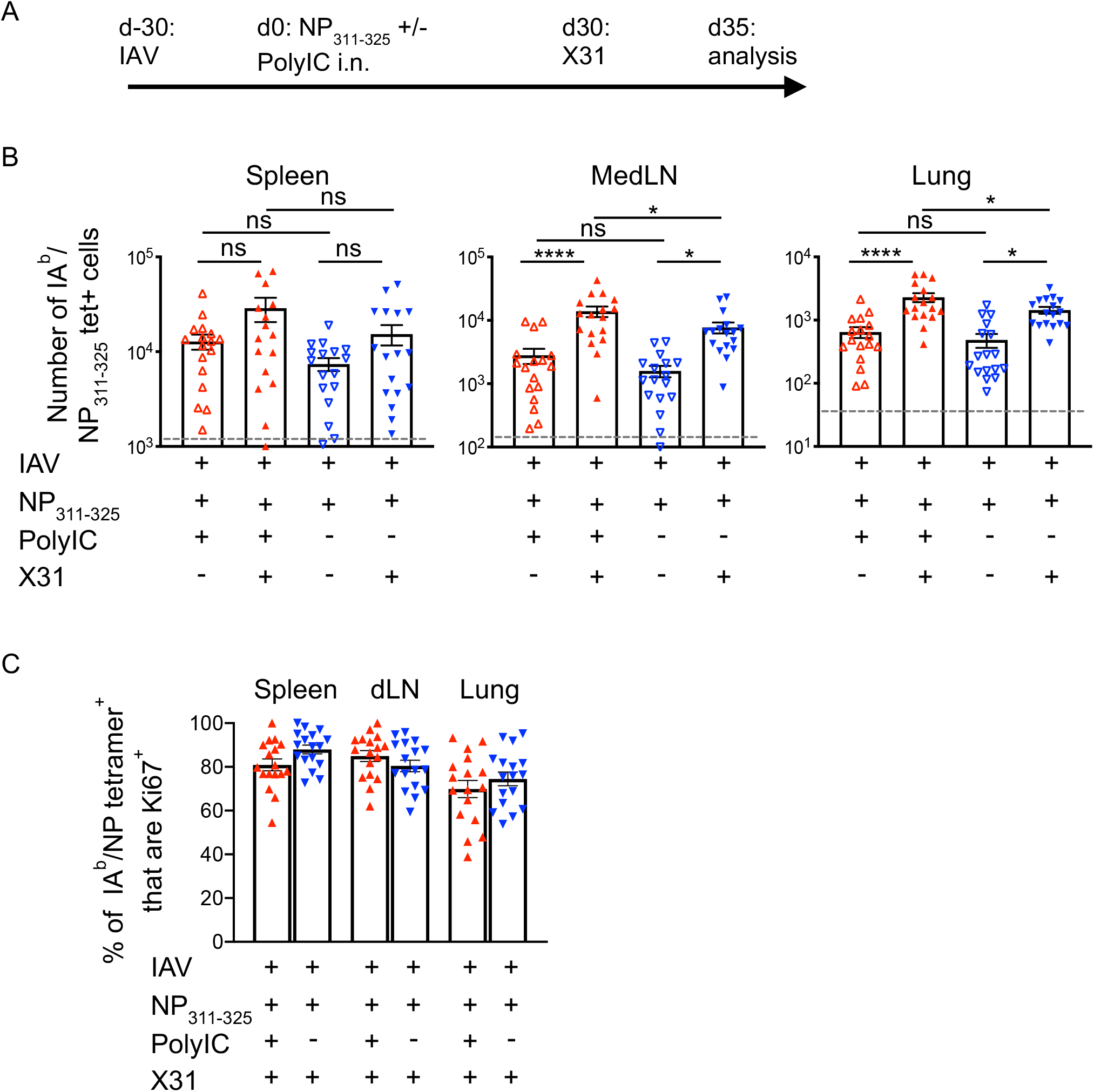
CD4 T cells exposed to peptide in the absence of adjuvant expand following re-infection with IAV. C57BL/6 mice were infected with IAV on day −30. On day 0, mice received NP_311-325_ +/− PolyIC i.n. and some of these mice were infected with 100PFU of X31 i.n. 30 days after this (A). The numbers of IA^b^/NP_311-325_ CD44^hi^ CD4 T cells were examined 5 days later in the spleen, MedLN, and lung (B) and their expression of Ki67 determined (C). In B and C, each symbol represents one mouse and error bars are SEM. In A, the grey dashed line represents the background staining in naïve animals. Data are combined from 3 experiments (5-6 mice/experiment). All statistics calculated using a one-way ANOVA with multiple comparisons; ns = not significant, * = <0.05, ** = <0.01, *** = <0.001, **** = <0.0001.

Five days following re-infection, we found significant increases in the numbers of antigen-specific CD4 T cells in the lungs and MedLN of mice regardless of their immunisation history (Figure 7B). This expansion was less clear in the spleen regardless of previous immunisation. In all organs, the majority of the antigen-specific CD4 T cells were Ki67 positive indicating a more robust response following IAV infection compared to immunisation (Figure 7C versus Figure 2C).

## Discussion

Memory CD4 T cells respond to low doses of antigen and costimulatory signals suggesting they will be refractory to tolerance induction(Blair et al., 2011, Holzer et al., 2003, London et al., 2000, MacLeod et al., 2006). This presents significant hurdles for therapies that aim to induce antigen-specific T cell tolerance(MacLeod and Anderton, 2015, Pearson et al., 2017, Ten Brinke et al., 2019).

Our previous(David et al., 2014) and current data demonstrate that some but not all functions of memory CD4 T cells are altered following the exposure of memory CD4 T cells to antigen delivered in the absence of adjuvant. Our findings demonstrate that tolerance induction in these cells is both subtle and complex. This contrasts with investigations of tolerance induction in naïve CD4 T cells that consistently show silencing of multiple T cell functions(Greenwald et al., 2005, Liu et al., 2019, Miller et al., 2007, Nurieva et al., 2006, Nurieva et al., 2011).

Consistently in our own and others research, memory CD4 T cells reactivated with antigen alone fail to accumulate following tertiary activation with antigen and adjuvant(David et al., 2014, Mackenzie et al., 2014). Our analysis of cell proliferation and survival signals suggest that this is not a consequence of reduced entry into the cell cycle nor low expression of anti-apoptosis molecules. Instead, our data indicate that memory CD4 T cells reactivated following exposure to antigen fail to complete mitosis, a characteristic associated with the phenomenon of mitotic catastrophe.

Mitotic catastrophe has mainly been studied in tumour cells treated with ionising radiation or drugs that cause DNA damage(Kimura et al., 2013, Maskey et al., 2013, Mc Gee, 2015, Vakifahmetoglu et al., 2008). Our transcriptomic data indicate that reactivated memory CD4 T cells previously exposed to tolerogenic signals have reduced expression of molecules involved in sensing and repairing DNA damage and in the control of various stages of the cell cycle. Our data suggest, therefore, a novel form of cell death for memory CD4 T cells.

In contrast to the poor accumulation of memory CD4 T cells exposed to tolerogenic signals, this did not shut down cytokine production. We found only small increases in cytokine-producing CD4 T cells following reactivation regardless of T cell activation history. If these cells are not undergoing cell division, they would not be at risk of death via mitotic catastrophe. These data suggest that non-cytokine producing memory CD4 T cells are more likely to proliferate than those committed to cytokine production. This agrees with the general concepts within the Tcentral/Teffector memory cell classification and with findings from ourselves and others that cells either making IFN-γ, or that are CD62L^lo^, proliferate poorly on reactivation(Dutta et al., 2013, MacLeod et al., 2008, Thomas et al., 2010).

We did find that memory CD4 T cells previously exposed to tolerogenic signals were unable to provide accelerated help for primary responding B cells, suggesting cell proliferation may be required for this functional response. This contrasts with our previous study in which memory cells exposed to tolerogenic signals could help primary responding B cells produce class switched antibody(David et al., 2014). There are multiple differences in experimental procedure between our previous study and the experiments here including the antigen (3K peptide versus NP_311-325_) form of priming (antigen versus infection) and route of tolerance induction (intravenous versus intranasal) that could explain this difference. Regardless, in our studies and similar research from others, memory CD4 T cells reactivated with antigen delivered without adjuvant accumulate poorly following a subsequent immunisation(David et al., 2014, Mackenzie et al., 2014). This suggests that consistently and, regardless of specificity, priming, memory cell location, and route of injection of tolerogenic signals, memory CD4 T cells reactivated with antigen in the absence of adjuvant proliferate poorly following reactivation with antigen and adjuvant.

Interestingly, we found that this poor accumulation could be rescued by re-infection with IAV, a much more potent challenge to the host than immunisation. These data suggest that while the responses of memory CD4 T cells can be moderated by exposure to tolerogenic signals, these cells may not be permanently silenced. An alternative explanation is that a portion of the memory CD4 T cells are not reactivated by antigen immunisations and therefore remain blind to the tolerogenic signals and free to respond to the infection. Teasing apart these two possibilities will require detailed understanding of the micro-location of memory CD4 T cells within peripheral and lymphoid organs and which antigen presenting cells reactivate memory CD4 T cells following immunisation and infection. A deeper understanding of these factors will be critical to address the most effective methods of antigen-specific tolerance strategies. Our data, moreover, demonstrate the importance of analysing multiple phenotypic and functional parameters in trials of antigen-specific therapy(Pearson et al., 2017, Ten Brinke et al., 2019).

## Materials and Methods

### Animals

To generate mice in which rtTA reports IL-2 expression we used recombineering to extract the upstream 8.389kb section of the IL-2 promoter from a Bacterial Artificial Chromosome (BAC) RP24208L3 (BAC resource at Children’s Hospital Oakland Research Institute, Buffalo, New York). This was subcloned into a plasmid containing the human CD2 locus control region and linked to the rtTA sequence. The transgene, cut and purified from the construct backbone, was used to create transgenic mice at the Transgenic mouse facility at National Jewish Health in Jackson Lab C57BL/6 animals. Two founder pups were identified by PCR but only one was fertile. Progeny of this animal were bred with B6.Cg- Tg(tetO-cre)1Jaw/J (006234) and B6.129X1-Gt(ROSA)26Sor^tm1(EYFP+)Cos^ (006148) both from Jackson Laboratories. 10 week old female C57BL/6 mice were purchased from Envigo (UK). TRACE and C57BL/6 mice were maintained at the University of Glasgow under standard animal husbandry conditions in accordance with UK home office regulations (Project License P2F28B003) and approved by the local ethics committee.

### Immunisations and infections

TRACE mice were given Dox+ chow (Envigo) for a total of 7 days starting two days prior to immunisation with 40μg of ovalbumin (OVA) protein (Worthington) conjugated to 20μm polyethylene carboxylate beads (Polysciences Inc.) with 20μg of polyinosinic:polycytidinic acid (InvivoGen) and 20μg of anti-CD40 (BioXcell) s.c in the scruff. Recipients of TRACE EYFP+ T cells were given 40μg of OVA with/out 10μg lipopolysaccharide i.v. in 100μl of PBS. NP_311-325_ was conjugated to OVA using Imject Maleimide-activated OVA according to the manufacturer’s instruction (ThermoFisher) and mice immunised with 1μg NP_311-325_-OVA delivered i.p with 0.1mg alum. For mitosis analysis, C57BL/6 mice were immunised with 20μg NP_311-325_ peptide (IDT) with 10μg of LPS i.v. After 30 days, they were re-immunised with 20μg NP_311-325_ peptide with/out 10μg LPS followed 30 days later by i.p. immunisation with 5μg NP-OVA with 0.1mg of alum i.p. For IAV studies, C57BL/6 mice were briefly anesthetised using inhaled isoflurane and infected with 200-300 plaque forming units of IAV strain WSN in 20μl of PBS intranasally (i.n.). IAV was prepared and titered in MDCK cells. Infected mice were rechallenged with 100PFU of X31 (kindly provided by Prof James Stewart, University of Liverpool). Infected mice were weighed daily. Any animals that lost more than 20% of their starting weight were humanely euthanised.

### FACS sorting and cell transfers

TRACE mice were euthanised 8 days post-immunisation and single cell suspensions were prepared. Lymphoid organs, including spleen, mediastinal, axillary, brachial and mesenteric lymph nodes, from individual mice were pooled and pre-enriched for CD4 T cells using EasySep™ Mouse T Cell Isolation Kit (Stemcell Technologies). Live, single, EYFP+ CD4 T cells negative for MHCII, B220, CD8 and F4/80 were sorted on a BD FACS Aria. Sorted cells were washed in PBS and 100,000 cells transferred i.v. into naïve C57BL/6 mice.

### Tissue preparation

Mice were euthanized either by cervical dislocation or with a rising concentration of carbon dioxide and perfused with PBS-5mM EDTA in experiments examining lungs. Spleen and mediastinal lymph nodes were processed by mechanical disruption. Single cell suspensions of lungs were prepared by digestion with 1mg/ml collagenase and DNAse (Sigma) for 40 minutes at 37°C. Red blood cells were lysed from spleen and lungs using lysis buffer (ThermoFisher).

### Flow cytometry

Single cell suspension were stained with PE-labeled IA^b^/NP_311-325_ (NIH tetramer core) at 37°C, 5% CO_2_ for 2 hours in complete RMPI (RPMI with 10% foetal calf serum, 100μg/ml penicillin-streptomycin and 2mM L-glutamine) containing Fc block (24G2). Surface antibodies were added and the cells incubated for a further 20minutes at 4°C. Antibodies used were: anti-CD4 BUV805 (BD Biosciences; clone: RM4-5) or CD4 APC-Alexa Fluor 780 (eBioscience; RM4-5), anti-CD44 BUV395 (BD Biosciences; clone: IM7), anti-CXCR5 BV785 (BioLegend; clone:L138D7), anti-PD-1 BV711 (BioLegend: 29F.1A12) and ‘dump’ antibodies: B220 (RA3-6B2), anti-CD8 (53-6.7) and MHC II (M5114) all on eFluor-450 (eBioscience). Cells were stained with a fixable viability dye eFluor 506 (eBioscience). In some cases, cells were then fixed with FoxP3 Transcription Factor Fixative kit (Thermofisher UK) and stained with anti-FoxP3 PeCy7 (eBioscience; FJK-16S), anti-Bcl2 FITC (Biolegend; Blc/10C4), anti-Ki67 BV605 (Biolegend; 16A8). Phosphorylated H3 was detected in cells fixed with 2%PFA/0.5% saponin using Alexa647-labelled anti-Histone H3 (pS28) (HTA28). Cells were acquired on a BD LSR or Fortessa and analysed using FlowJo (version 10 Treestar).

### T cell cytokine analysis

Bone marrow derived dendritic cells were cultured as described(Inaba et al., 1992) in complete RPMI supplemented with X-63 supernatant for 7 days. A single cell suspension was incubated with 10μg/ml NP_311-325_ peptide for 2 hours prior to co-culture with lungs, spleen or lymph node cells in complete RMPI at a ratio of approximately 10 T cells to 1 DC in the presence of Golgi Plug (BD Bioscience). Co-cultures were incubated at 37°C, 5% CO_2_ for 6 hours. Cells were incubated with Fc block and surface stained with anti-CD4 BUV805 (BD Biosciences; clone: RM4-5) or CD4 APC-Alexa Fluor 780 (eBioscience; RM4-5), anti-CD44 BUV395 (BD Biosciences; clone: IM7) and ‘dump’ antibodies: B220 (clone: RA3-6B2), CD8 (53-6.7) and MHC II (clone: M5114) all on eFluor-450 (eBioscience). Cells were fixed with cytofix/cytoperm (BD Bioscience) for 20 minutes at 4°C and stained in permwash buffer with anti-cytokine antibodies for one hour at room temperature (anti-IFN-γ PE (XMG1.2;), anti-TNF Alexa-Fluor-488 (MP6-XT22) anti-IL-2 APC (JES6-5H4) all from eBioscience.

### RNA isolation for RNA-seq

CD4+ EYFP+ cells from the spleen, mediastinal, axillary, brachial and mesenteric lymph nodes of TRACE cell recipients were FACS sorted as above and cell pellets were stored at − 20°C prior to RNA extraction. RNA was extracted and purified from single cell suspensions using RNeasy Micro Kit (Qiagen) according to manufacturer’s instructions.

### RNA analysis

Sequencing and library prep were conducted by LCSciences Ltd. Total RNA was extracted using Trizol reagent (Invitrogen, CA, USA). Total RNA quantity and purity were analysed using a Bioanalyzer 2100 and RNA 6000 Nano LabChip Kit (Agilent, CA, USA), all samples had RIN numbers >7.0. Approximately 10μg of total RNA was subjected to isolate Poly (A) mRNA with poly-T oligoattached magnetic beads (Invitrogen). Following purification, the poly(A)- or poly(A)+ RNA fractions were fragmented using divalent cations under elevated temperature. The cleaved RNA fragments were reverse-transcribed to create the final cDNA library in accordance with the protocol for the mRNA-Seq sample preparation kit (Illumina, San Diego, USA). The average insert size for the paired-end libraries was 300 bp (±50 bp). Paired-end sequencing was done on an Illumina Hiseq 4000 (lc-bio, China). Cutadapt software(Martin, 2011) was used to remove low quality reads and adaptor sequences. High quality reads were then mapped to a C57BL/6 mouse reference genome to which the EYFP sequence was added using HISAT2(Kim et al., 2015). Readcounts were obtained from bam files with FeatureCounts using the default parameters. Differential expressed genes (DEGs) were obtained with DESeq2, using RStudio (RStudio Inc). DEGs were visualised with a volcano plot using the ‘Enhancedvolcano’ package within R. DEGs with a fold change of at least 3 and q-values less than p=0.05 were classed as statistically significant. Heatmaps were generated using the heatmap2 function, using the DESeq2 normalised counts and Panther(Mi et al., 2019) used to determine GO biological processes. GSEA analysis was conducted using UC San Diego/Broad Institute’s GSEA software(Mootha et al., 2003, Subramanian et al., 2005).

### ELISA

OVA specific antibody ELISAs were carried out as descried(David et al., 2014). Serum from immunised and control mice was titrated in 2-fold serial dilution on plates coated with OVA protein. Anti-mouse IgG, IgG1 or IgG2c biotin detection antibodies (Thermofisher, UK) were used with Extravidin-peroxidase (Sigma Aldrich) and SureBlueTMB substrate (KPL). Absorbance was measured at 450nm using a Sunrise Absorbance Reader (Tecan). The absorbance of each sample was normalised to a positive control on each plate after the background absorbance from a blank well had been removed.

### Statistical analysis

Data were analysed using Prism version 7 software (GraphPad). Differences between groups were analysed by unpaired ANOVAs or T-tests as indicated in figure legends. In all figures * represents a p value of <0.05; **: p>0.01, ***: p>0.001, ****: p>0.0001.

## Acknowledgements

We thank the staff within the Institute of Infection, Immunity and Inflammation Flow Cytometry Facility, the Joint and Central Research Facility at the University of Glasgow for technical assistance. We thank the NIH tetramer core facility for provision of IA^b^-NP_311-325_-PE and Prof James Stewart (University of Liverpool) for IAV X31. We thank Dr Claire McIntyre and Mr Colin Chapman for technical assistance and Dr Catharien Hilkens (University of Newcastle) and Prof David Withers (University of Birmingham) for critical reading of the manuscript. This work was supported by an Arthritis Research UK Career Development Fellowship (19905) and a Marie Curie Fellowship (334430) to MKLM, by a DTP-MRC studentship to JIG (MR/JR50032X/1) and by the Howard Hughes Medical Institute. Transcriptomics data have been deposited on GEO, accession number GSE145310.

## Author Contribution

JIG designed and performed experiments, analysed data and wrote the manuscript; TO and S-AK analysed data; FM, ETC, LG, JLM, JWK and PM designed and produced essential tools; PG designed experiments; MKLM designed and performed the research, analysed data, and wrote the manuscript. All authors approved the manuscript.

## Supplementary Figures

**Supplementary Figure 1:**
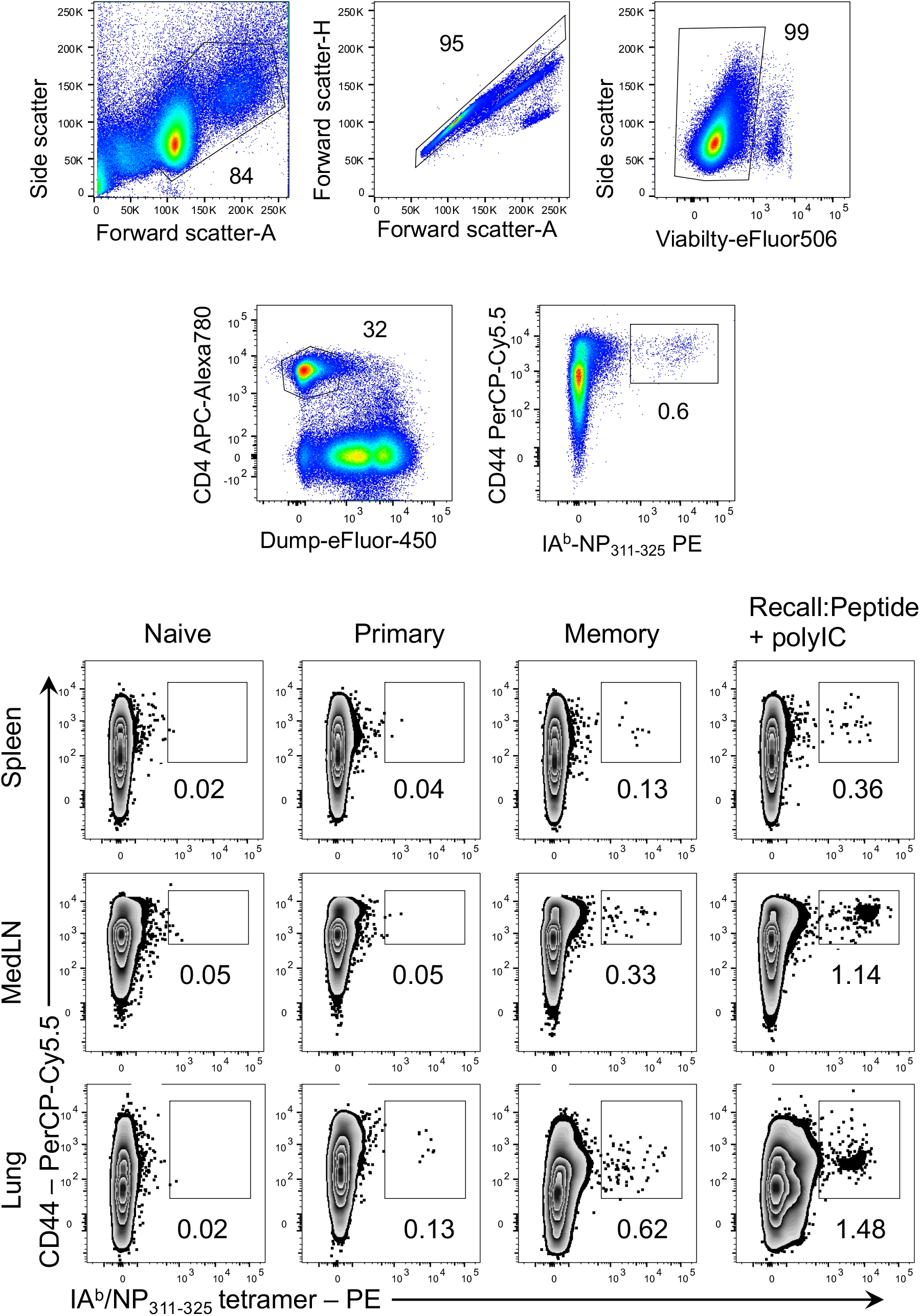
Gating strategy for IA^b^/NP_311-325_ tetramer+ CD44^hi^ cells. C57BL/6 mice were infected with IAV i.n. and some were given NP_311-325_ peptide+PolyIC i.n. on day 30. The percentages of IA^b^/NP_311-325_ tetramer+ CD44^hi^ CD4 T cells examined 35 days later in spleen, mediastinal LN, and lung. Cells are gated on live CD4+ lymphocytes that are negative for B220, F4/80, CD8, and MHCII+ as shown in the gating strategy. The numbers on the graph show the percentages of the cells present in the gate within each plot.

**Supplementary Figure 2:**
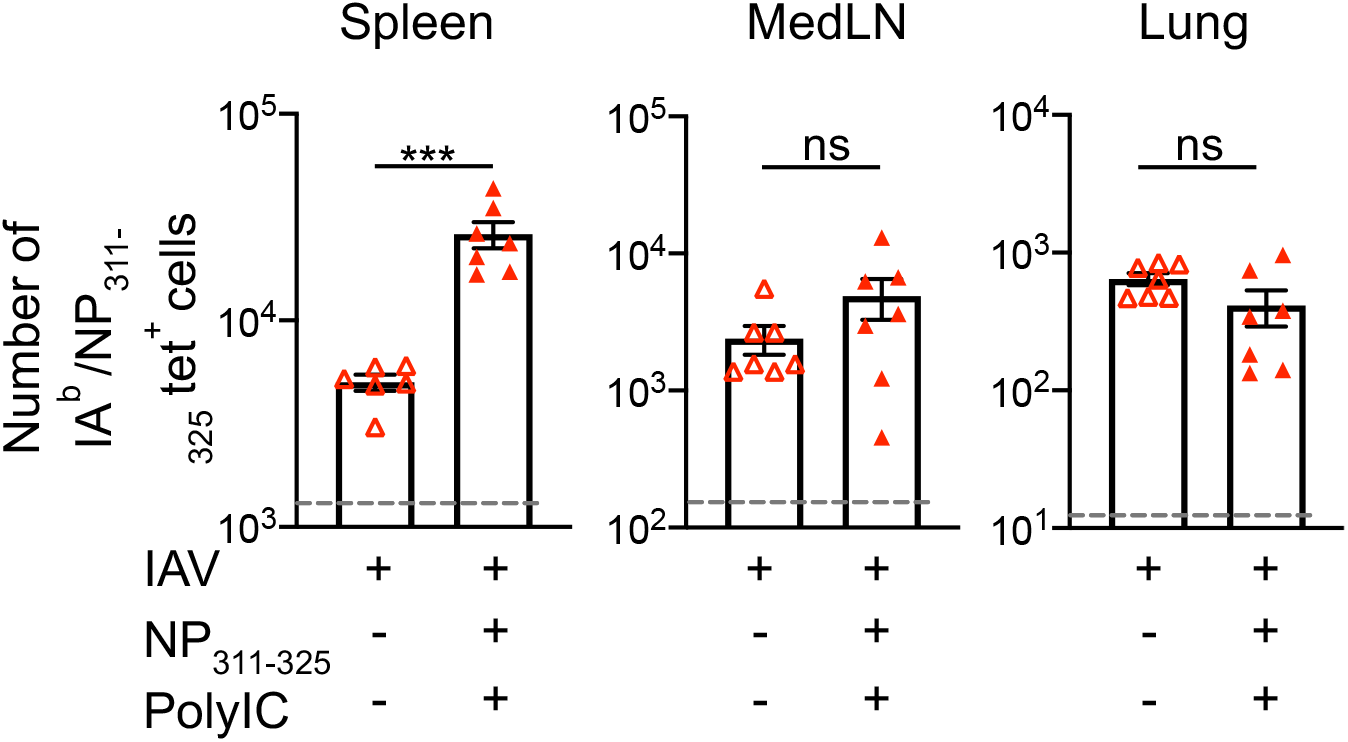
Lung memory NP_311-325_-antigen-specific CD4 T cells are not reactivated by antigen and adjuvant delivered i.v. C57BL/6 mice were infected with IAV i.n. on day −30. On day 0, some of these mice were immunised with NP_311-325_, + PolyIC intravenously. The numbers of IA^b^/NP_311-325_ CD44^hi^ CD4 T cells were examined 5 days later in the spleen, MedLN, and lung. Each symbol represents one mouse and error bars are SEM. The grey dashed line represents the background staining in naïve animals. Data are combined from two experiments (3-4mice/experiment). Statistics calculated using a one-way ANOVA with multiple comparisons; ns = not significant, * = <0.05, ** = <0.01, *** = <0.001, **** = <0.0001.

**Supplementary Figure 3:**
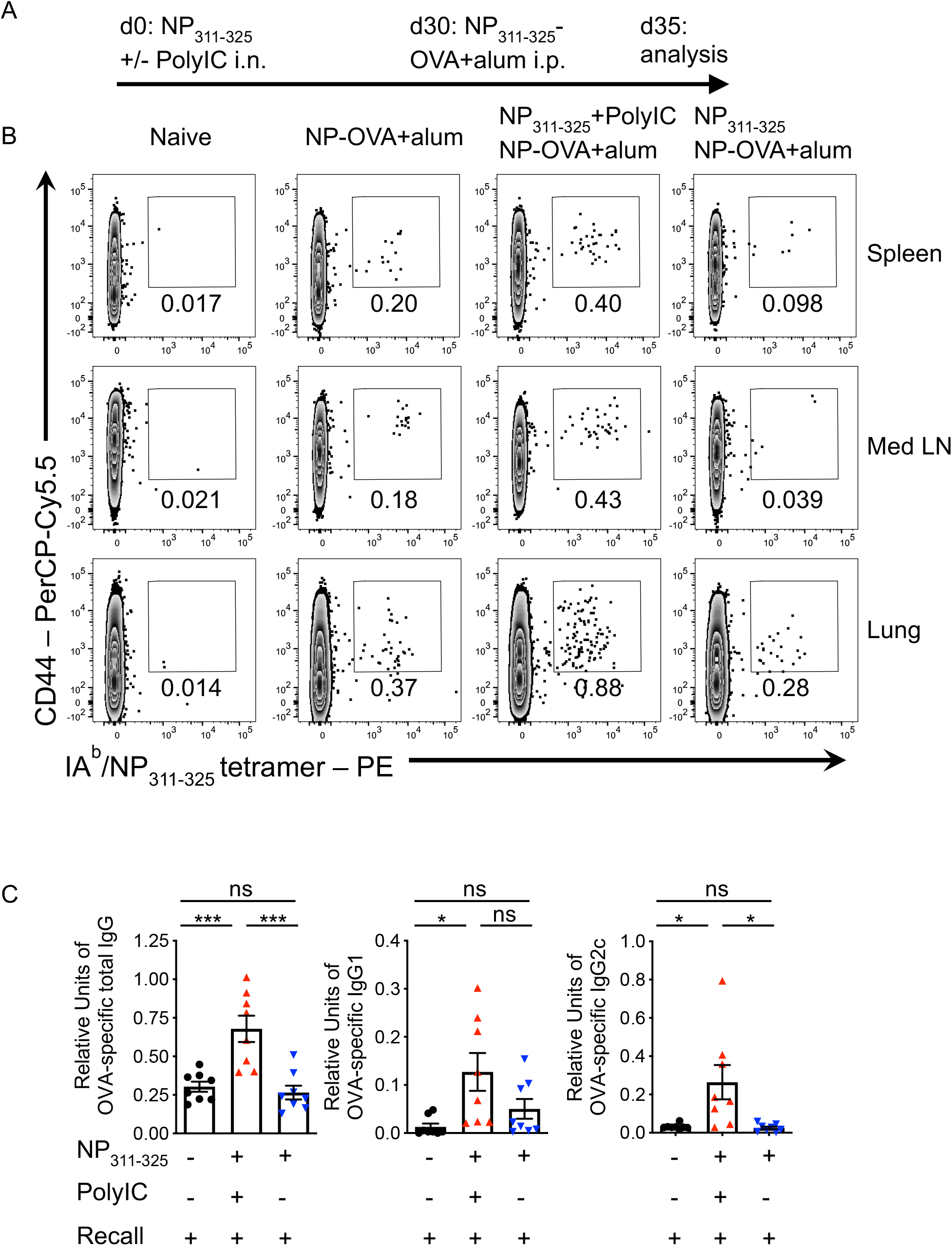
Instillation of NP_311-325_ peptide induces functional tolerance in naive animals. C57BL/6 mice were instilled with NP_311-325_ peptide +/− PolyIC on day 0 and immunised with NP_311-325_-OVA and alum i.p. 30 days later (A). The percentages of IA^b^/NP_311-325_ CD44^hi^ CD4 T cells were examined 5 days after the recall immunisation (B). Cells are gated on live CD4+ lymphocytes that are negative for B220, CD8, F4/80, and MHCII+. The numbers on the graph show the percentages of the cells present in the gate within each plot. The levels of IgG, IgG1 and IgG2c anti-OVA antibodies was determined on day 5 in the serum (C). Data are from 2 experiments with 4mice/group. In C each point represents one mouse and the error bars are SEM). All statistics calculated using a one-way ANOVA with multiple comparisons; ns = not significant, * = <0.05, ** = <0.01, *** = <0.001.

**Supplementary Figure 4:**
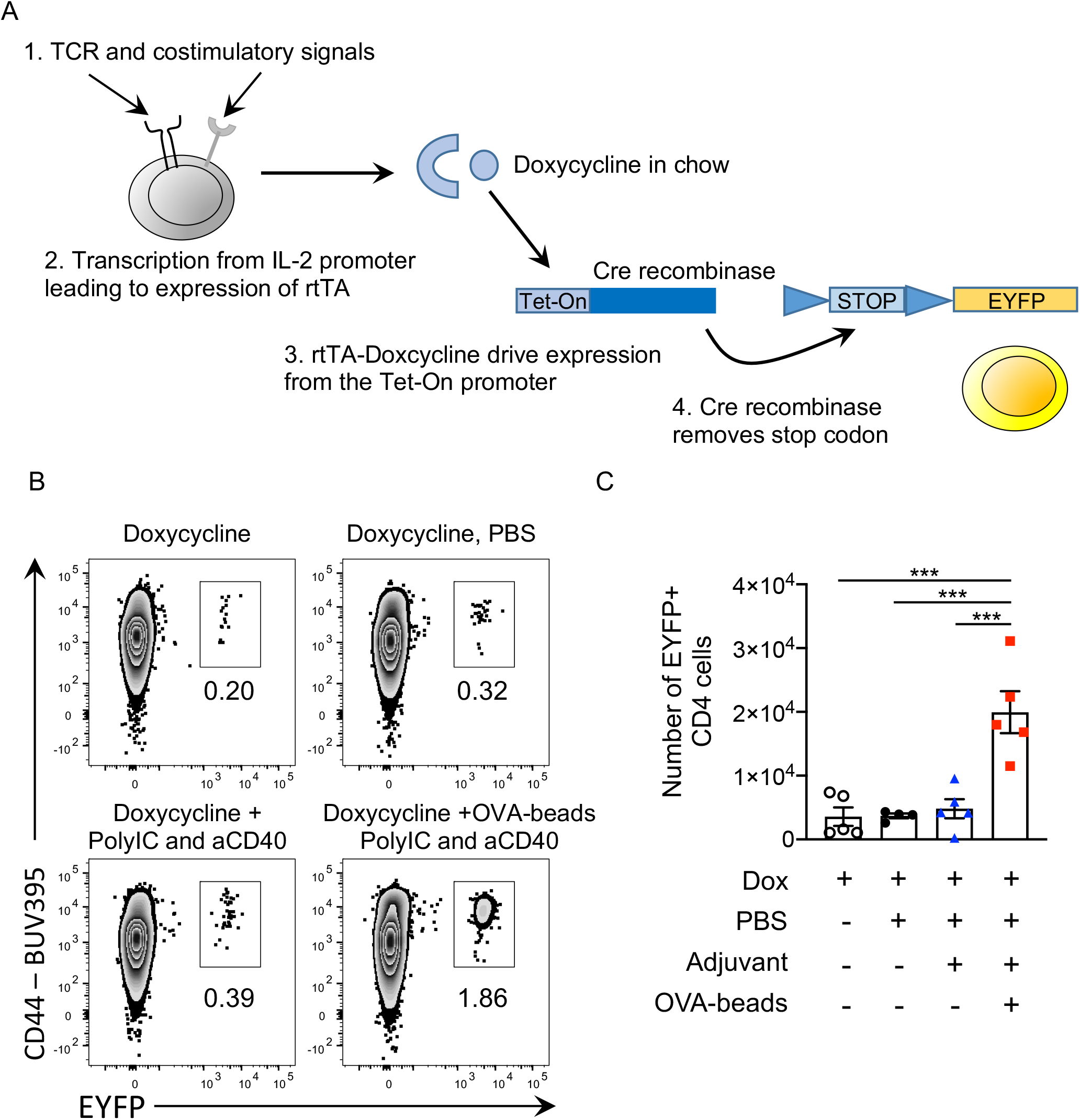
TRACE mice enable identification of antigen-reactive CD4 T cells. In TRACE mice, activation through the TCR in the presence of costimulatory signals leads to the expression of rtTA driven by the interleukin 2 promoter. Only in the presence of doxycycline will rtTA be able to bind to the tet-ON promoter leading to the expression of Cre recombinase. Cre recombinase removes the stop codon at the Rosa locus allowing permanent expression of EYFP (A). In B, TRACE mice were given doxycycline chow from day minus 2 until day 5 and injected in the s.c. in the scruff on day 0 with nothing, PBS, anti-CD40 and PolyIC, or OVA protein conjugated to 20μm beads delivered with anti-CD40 and PolyIC. On day 9, the brachial and axillary lymph nodes were examined by flow cytometry. Cells are gated on live lymphocytes that were CD4+, and negative for B220, F4/80, MHCII and CD8. The numbers show the percentage of CD4+ cells that are in the indicated gates (A). In B, the numbers of EYFP+ CD4 T cells in the lymph nodes are shown with each symbol representing one mouse. Statistics calculated using a one-way ANOVA with multiple comparisons; *** = <0.001.

**Supplementary Figure 5:**
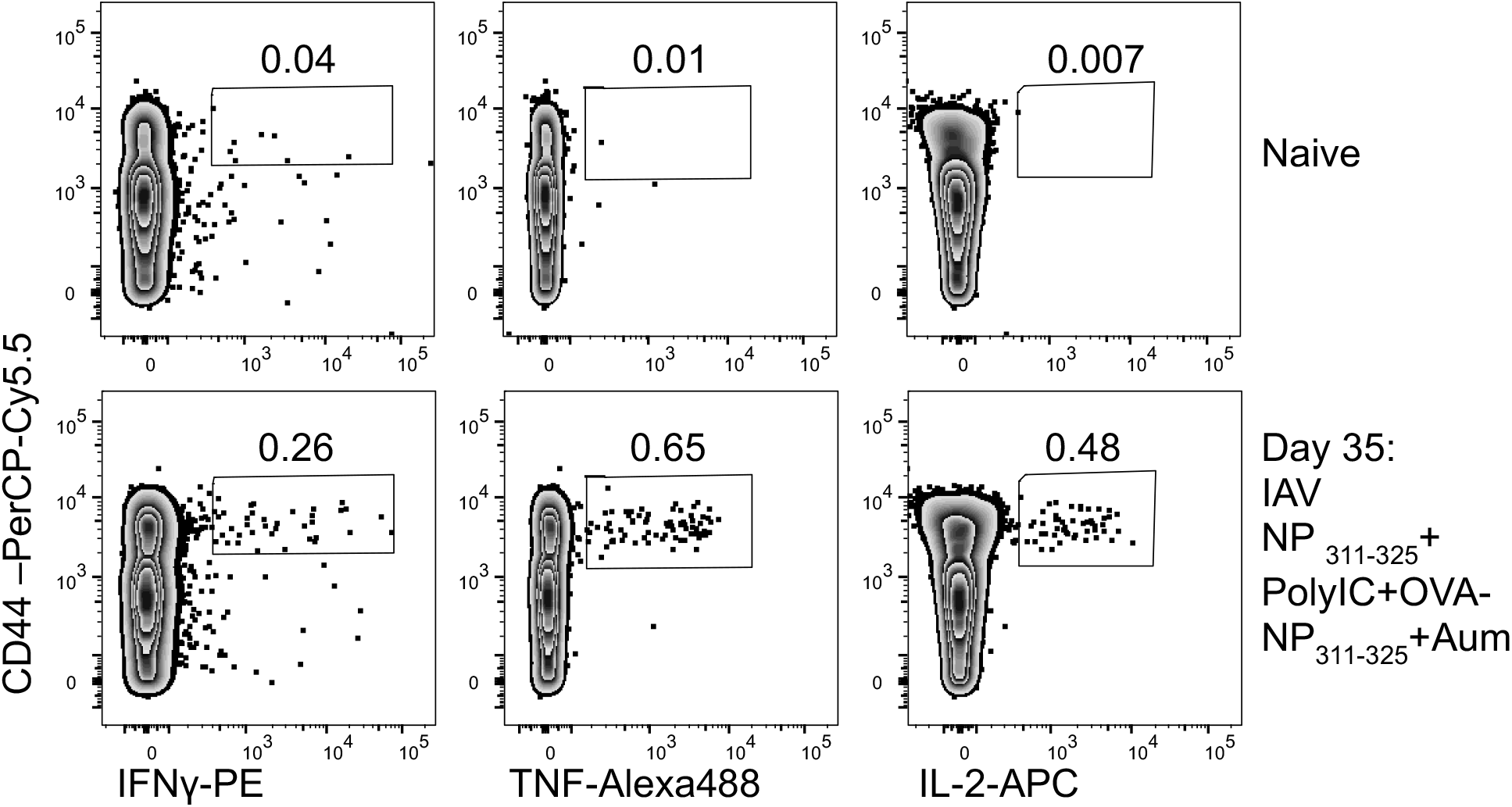
Identification of NP_311-325_ specific cytokine producing CD4 T cells. C57BL/6 mice were infected with IAV on day −30. On day 0, mice received NP_311-325_ in the +/− PolyIC and some of these mice immunised i.p with NP-OVA with alum on day 30. On day 35, cells from the spleen were co-cultured with bmDCs loaded with NP_311-325_ for 6 hours in the presence of Golgi Plug and the percentages of IFN-γ, TNF and IL-2 producing CD44^hi^ CD4+ T cells examined. Cells are gated on live CD4+ lymphocytes as in gating strategy in SF1. Data are representative of 3 experiments (3-5mice/experiment).

## References

Blair, D. A., Turner, D. L., Bose, T. O., Pham, Q. M., Bouchard, K. R., Williams, K. J., Mcaleer, J. P., Cauley, L. S., Vella, A. T. & Lefrancois, L. 2011. Duration of antigen availability influences the expansion and memory differentiation of T cells. J Immunol, 187, 2310–21.

Bohmer, R. M., Bandala-Sanchez, E. & Harrison, L. C. 2011. Forward light scatter is a simple measure of T-cell activation and proliferation but is not universally suited for doublet discrimination. Cytometry A, 79, 646–52.

Cope, A. P., Schulze-Koops, H. & Aringer, M. 2007. The central role of T cells in rheumatoid arthritis. Clin Exp Rheumatol, 25, S4–11.

David, A., Crawford, F., Garside, P., Kappler, J. W., Marrack, P. & Macleod, M. 2014. Tolerance induction in memory CD4 T cells requires two rounds of antigen-specific activation. Proc Natl Acad Sci U S A, 111, 7735–40.

Dutta, A., Miaw, S. C., Yu, J. S., Chen, T. C., Lin, C. Y., Lin, Y. C., Chang, C. S., He, Y. C., Chuang, S. H., Yen, M. I. & Huang, C. T. 2013. Altered T-bet dominance in IFN-gamma-decoupled CD4+ T cells with attenuated cytokine storm and preserved memory in influenza. J Immunol, 190, 4205–14.

Greenwald, R. J., Freeman, G. J. & Sharpe, A. H. 2005. The B7 family revisited. Annu Rev Immunol, 23, 515–48.

Gunawardana, N. C. & Durham, S. R. 2018. New approaches to allergen immunotherapy. Ann Allergy Asthma Immunol, 121, 293–305.

Hans, F. & Dimitrov, S. 2001. Histone H3 phosphorylation and cell division. Oncogene, 20, 3021–7.

Hardwick, K. G. & Murray, A. W. 1995. Mad1p, a phosphoprotein component of the spindle assembly checkpoint in budding yeast. J Cell Biol, 131, 709–20.

Hartigan, C. R., Sun, H. & Ford, M. L. 2019. Memory T-cell exhaustion and tolerance in transplantation. Immunol Rev, 292, 225–242.

Holzer, U., Kwok, W. W., Nepom, G. T. & Buckner, J. H. 2003. Differential antigen sensitivity and costimulatory requirements in human Th1 and Th2 antigen-specific CD4+ cells with similar TCR avidity. J Immunol, 170, 1218–23.

Inaba, K., Inaba, M., Romani, N., Aya, H., Deguchi, M., Ikehara, S., Muramatsu, S. & Steinman, R. M. 1992. Generation of large numbers of dendritic cells from mouse bone marrow cultures supplemented with granulocyte/macrophage colony-stimulating factor. J Exp Med, 176, 1693–702.

Jaigirdar, S. A. & Macleod, M. K. 2015. Development and Function of Protective and Pathologic Memory CD4 T Cells. Front Immunol, 6, 456.

Jenkins, M. K. & Schwartz, R. H. 1987. Antigen presentation by chemically modified splenocytes induces antigen-specific T cell unresponsiveness in vitro and in vivo. J Exp Med, 165, 302–19.

Joukov, V. & De Nicolo, A. 2018. Aurora-PLK1 cascades as key signaling modules in the regulation of mitosis. Sci Signal, 11.

Kim, D., Langmead, B. & Salzberg, S. L. 2015. HISAT: a fast spliced aligner with low memory requirements. Nat Methods, 12, 357–60.

Kimura, M., Yoshioka, T., Saio, M., Banno, Y., Nagaoka, H. & Okano, Y. 2013. Mitotic catastrophe and cell death induced by depletion of centrosomal proteins. Cell Death Dis, 4, e603.

Kurche, J. S., Burchill, M. A., Sanchez, P. J., Haluszczak, C. & Kedl, R. M. 2010. Comparison of OX40 ligand and CD70 in the promotion of CD4+ T cell responses. J Immunol, 185, 2106–15.

Liblau, R. S., Tisch, R., Shokat, K., Yang, X., Dumont, N., Goodnow, C. C. & Mcdevitt, H. O. 1996. Intravenous injection of soluble antigen induces thymic and peripheral T-cells apoptosis. Proc Natl Acad Sci U S A, 93, 3031–6.

Liu, X., Wang, Y., Lu, H., Li, J., Yan, X., Xiao, M., Hao, J., Alekseev, A., Khong, H., Chen, T., Huang, R., Wu, J., Zhao, Q., Wu, Q., Xu, S., Wang, X., Jin, W., Yu, S., Wang, Y., Wei, L., Wang, A., Zhong, B., Ni, L., Liu, X., Nurieva, R., Ye, L., Tian, Q., Bian, X. W. & Dong, C. 2019. Genome-wide analysis identifies NR4A1 as a key mediator of T cell dysfunction. Nature, 567, 525–529.

Lizarraga, S. B., Margossian, S. P., Harris, M. H., Campagna, D. R., Han, A. P., Blevins, S., Mudbhary, R., Barker, J. E., Walsh, C. A. & Fleming, M. D. 2010. Cdk5rap2 regulates centrosome function and chromosome segregation in neuronal progenitors. Development, 137, 1907–17.

London, C. A., Lodge, M. P. & Abbas, A. K. 2000. Functional responses and costimulator dependence of memory CD4+ T cells. J Immunol, 164, 265–72.

Mackenzie, K. J., Nowakowska, D. J., Leech, M. D., Mcfarlane, A. J., Wilson, C., Fitch, P. M., O’connor, R. A., Howie, S. E., Schwarze, J. & Anderton, S. M. 2014. Effector and central memory T helper 2 cells respond differently to peptide immunotherapy. Proc Natl Acad Sci U S A, 111, E784–93.

Macleod, M., Kwakkenbos, M. J., Crawford, A., Brown, S., Stockinger, B., Schepers, K., Schumacher, T. & Gray, D. 2006. CD4 memory T cells survive and proliferate but fail to differentiate in the absence of CD40. J Exp Med, 203, 897–906.

Macleod, M. K. & Anderton, S. M. 2015. Antigen-based immunotherapy (AIT) for autoimmune and allergic disease. Curr Opin Pharmacol, 23, 11–6.

Macleod, M. K., David, A., Jin, N., Noges, L., Wang, J., Kappler, J. W. & Marrack, P. 2013. Influenza nucleoprotein delivered with aluminium salts protects mice from an influenza A virus that expresses an altered nucleoprotein sequence. PLoS One, 8, e61775.

Macleod, M. K., David, A., Mckee, A. S., Crawford, F., Kappler, J. W. & Marrack, P. 2011. Memory CD4 T cells that express CXCR5 provide accelerated help to B cells. J Immunol, 186, 2889–96.

Macleod, M. K., Mckee, A., Crawford, F., White, J., Kappler, J. & Marrack, P. 2008. CD4 memory T cells divide poorly in response to antigen because of their cytokine profile. Proc Natl Acad Sci U S A, 105, 14521–6.

Martin, M. 2011. Cutadapt removes adapter sequences from high-throughput sequencing reads. EMBnet.journal, 17.

Maskey, D., Yousefi, S., Schmid, I., Zlobec, I., Perren, A., Friis, R. & Simon, H. U. 2013. ATG5 is induced by DNA-damaging agents and promotes mitotic catastrophe independent of autophagy. Nat Commun, 4, 2130.

Mc Gee, M. M. 2015. Targeting the Mitotic Catastrophe Signaling Pathway in Cancer. Mediators Inflamm, 2015, 146282.

Mcginley, A. M., Edwards, S. C., Raverdeau, M. & Mills, K. H. G. 2018. Th17cells, gammadelta T cells and their interplay in EAE and multiple sclerosis. J Autoimmun.

Mi, H., Muruganujan, A., Ebert, D., Huang, X. & Thomas, P. D. 2019. PANTHER version 14: more genomes, a new PANTHER GO-slim and improvements in enrichment analysis tools. Nucleic Acids Res, 47, D419–D426.

Miller, S. D., Turley, D. M. & Podojil, J. R. 2007. Antigen-specific tolerance strategies for the prevention and treatment of autoimmune disease. Nat Rev Immunol, 7, 665–77.

Mootha, V. K., Lindgren, C. M., Eriksson, K. F., Subramanian, A., Sihag, S., Lehar, J., Puigserver, P., Carlsson, E., Ridderstrale, M., Laurila, E., Houstis, N., Daly, M. J., Patterson, N., Mesirov, J. P., Golub, T. R., Tamayo, P., Spiegelman, B., Lander, E. S., Hirschhorn, J. N., Altshuler, D. & Groop, L. C. 2003. PGC-1alpha-responsive genes involved in oxidative phosphorylation are coordinately downregulated in human diabetes. Nat Genet, 34, 267–73.

Musacchio, A. 2015. The Molecular Biology of Spindle Assembly Checkpoint Signaling Dynamics. Curr Biol, 25, R1002–18.

Nitta, M., Kobayashi, O., Honda, S., Hirota, T., Kuninaka, S., Marumoto, T., Ushio, Y. & Saya, H. 2004. Spindle checkpoint function is required for mitotic catastrophe induced by DNA-damaging agents. Oncogene, 23, 6548–58.

Nurieva, R., Thomas, S., Nguyen, T., Martin-Orozco, N., Wang, Y., Kaja, M. K., Yu, X. Z. & Dong, C. 2006. T-cell tolerance or function is determined by combinatorial costimulatory signals. EMBO J, 25, 2623–33.

Nurieva, R. I., Liu, X. & Dong, C. 2011. Molecular mechanisms of T-cell tolerance. Immunol Rev, 241, 133–44.

Pearson, R. M., Casey, L. M., Hughes, K. R., Miller, S. D. & Shea, L. D. 2017. In vivo reprogramming of immune cells: Technologies for induction of antigen-specific tolerance. Adv Drug Deliv Rev, 114, 240–255.

Raphael, I., Joern, R. R. & Forsthuber, T. G. 2020. Memory CD4(+) T Cells in Immunity and Autoimmune Diseases. Cells, 9.

Rayner, F. & Isaacs, J. D. 2018. Therapeutic tolerance in autoimmune disease. Semin Arthritis Rheum, 48, 558–562.

Santamaria, D., Barriere, C., Cerqueira, A., Hunt, S., Tardy, C., Newton, K., Caceres, J. F., Dubus, P., Malumbres, M. & Barbacid, M. 2007. Cdk1 is sufficient to drive the mammalian cell cycle. Nature, 448, 811–5.

Serra, P. & Santamaria, P. 2019. Antigen-specific therapeutic approaches for autoimmunity. Nat Biotechnol, 37, 238–251.

Shang, Z. F., Huang, B., Xu, Q. Z., Zhang, S. M., Fan, R., Liu, X. D., Wang, Y. & Zhou, P. K. 2010. Inactivation of DNA-dependent protein kinase leads to spindle disruption and mitotic catastrophe with attenuated checkpoint protein 2 Phosphorylation in response to DNA damage. Cancer Res, 70, 3657–66.

Subramanian, A., Tamayo, P., Mootha, V. K., Mukherjee, S., Ebert, B. L., Gillette, M. A., Paulovich, A., Pomeroy, S. L., Golub, T. R., Lander, E. S. & Mesirov, J. P. 2005. Gene set enrichment analysis: a knowledge-based approach for interpreting genome-wide expression profiles. Proc Natl Acad Sci U S A, 102, 15545–50.

Ten Brinke, A., Martinez-Llordella, M., Cools, N., Hilkens, C. M. U., Van Ham, S. M., Sawitzki, B., Geissler, E. K., Lombardi, G., Trzonkowski, P. & Martinez-Caceres, E. 2019. Ways Forward for Tolerance-Inducing Cellular Therapies-an AFACTT Perspective. Front Immunol, 10, 181.

Thomas, P. G., Brown, S. A., Morris, M. Y., Yue, W., So, J., Reynolds, C., Webby, R. J. & Doherty, P. C. 2010. Physiological numbers of CD4+ T cells generate weak recall responses following influenza virus challenge. J Immunol, 184, 1721–7.

Vakifahmetoglu, H., Olsson, M. & Zhivotovsky, B. 2008. Death through a tragedy: mitotic catastrophe. Cell Death Differ, 15, 1153–62.

Zhang, J., Gao, W., Yang, X., Kang, J., Zhang, Y., Guo, Q., Hu, Y., Xia, G. & Kang, Y. 2013. Tolerogenic vaccination reduced effector memory CD4 T cells and induced effector memory Treg cells for type I diabetes treatment. PLoS One, 8, e70056.

Zhang, X., Liu, D., Lv, S., Wang, H., Zhong, X., Liu, B., Wang, B., Liao, J., Li, J., Pfeifer, G. P. & Xu, X. 2009. CDK5RAP2 is required for spindle checkpoint function. Cell Cycle, 8, 1206–16.

